# Poretools: a toolkit for analyzing nanopore sequence data

**DOI:** 10.1101/007401

**Authors:** Nicholas J. Loman, Aaron R. Quinlan

## Abstract

**Motivation:** Nanopore sequencing may be the next disruptive technology in genomics. Nanopore sequencing has many attractive properties including the ability to detect single DNA molecules without prior amplification, the lack of reliance on expensive optical components, and the ability to sequence very long fragments. The MinION from Oxford Nanopore Technologies (ONT) is the first nanopore sequencer to be commercialised and is now available to early-access users. The MinION™ is a USB-connected, portable nanopore sequencer which permits real-time analysis of streaming event data. A cloud-based service is available to translate events into nucleotide base calls. However, software support to deal with such data is limited, and the community lacks a standardized toolkit for the analysis of nanopore datasets.

**Results:** We introduce `poretools`, a flexible toolkit for manipulating and exploring datasets generated by nanopore sequencing devices from MinION for the purposes of quality control and downstream analysis. `Poretools` operates directly on the native FAST5 (a variant of the HDF5 standard) file format produced by ONT and provides a wealth of format conversion utilities and data exploration and visualization tools.

**Availability and implementation:** Poretools is open source software and is written in Python as both a suite of command line utilities and a Python application programming interface. Source code and user documentation are freely available in Github at https://github.com/arq5x/poretools

**Supplementary information:** An IPython notebook demonstrating the use and functionality of `poretools` in greater detail is available from the Github repository.

## 1 INTRODUCTION

The idea of using biological nanopores for DNA sequencing was proposed almost twenty years ago (Church *et al*. (1995); Kasianowicz *et al*. (1996)). This approach relies on the direct, electrical detection of single DNA strands in contact with an individual pore. Single molecule detection and the absence of a prior amplification step means that extremely long fragments can be sequenced without any loss in quality. In May 2014, Oxford Nanopore Technologies released MinION™, the first commercially-available nanopore DNA sequencing device. MinION™ is noteworthy for its portability, size (around the same length as an iPhone™) and USB 3.0 connectivity, meaning it can be run on a standard Internet-connected laptop. Laboratories throughout the world are actively evaluating this device for a broad range of applications. Sequencing with the MinION yields raw signals reflecting modulation of the ionic current at each pore by a DNA molecule. The resulting time-series of nanopore translocation, ‘events’, are base-called by proprietary software running as a cloud service. The resulting files for each sequenced read are stored in ‘FAST5’ format, a specialization of the HDF5 format (The HDF Group (1997)). This format is widely used in scientific computing applications. However, at present, no specific software is available to deal with downstream analysis starting with this file format.

## 2 FEATURES AND METHODS

We have developed `poretools`, an open source software toolkit that addresses the pressing need for methods to manipulate the FAST5 format and permit explorations of the raw nanopore event data and the resulting DNA sequences. `Poretools` provides an extensive set of data analysis methods that operate directly on either a single FAST5 file or a set of files from one or more sequencing runs. A Python programming library is provided to facilitate access to the FAST5 file structure and enable other researchers to extend the tools and create new analytical methods. In the following sections, we summarize and provide examples of the functionality currently available in `poretools`.

### 2.1 Format Conversion

The most fundamental requirement provided by `poretools` is the ability to convert the sequencing data resulting from a MinION run from HDF5/FAST5 format to either FASTA or FASTQ format in order to facilitate analyses with sequence alignment and/or assembly software. This is accomplished with the `fasta` and `fastq` commands in the `poretools` suite.

~~~
poretools **fastq** /path/to/fast5/example.fast5
poretools **fasta** /path/to/fast5/example.fast5
~~~

At the time of writing, each MinION run generates individual HDF5/FAST5 files for each sequenced read. Consequently, there are often tens of thousands of individuals files that must be stored for a single experiment. `Poretools` provides two different strategies for facilitating the analysis of such datasets.

The first approach allows one to execute a `poretools` command on an entire directory of FAST5 files.

~~~
poretools **fastq** /path/to/fast5/directory/
~~~

Alternatively, we provide a utility to combine a set of HDF5/FAST5 files into a single TAR file. This allows an entire run to be archived into one file and once combined, all other `poretools` commands are able to operate on each HDF5/FAST5 file therein. For example:

~~~
poretools combine -o run.tar /path/to/fast5/directory/
poretools **fastq** run.tar
~~~

### 2.2 Data exploration and visualization

Tthere is a need to visualize MinION™ run performance in order to assess its quality and troubleshoot different fragmentation and library preparation strategies. `Poretools` provides two utilties, `hist`, and `yield_plot` that characterize the fragment size distribution and display a collector’s curve of the overall sequencing yield respectively. Examples of the figures created by these utilities are shown (Figure 1A and 1B) and example commands are provided below.

~~~
poretools **hist** /path/to/fast5/directory/
poretools **yield_plot** /path/to/fast5/directory/
~~~

**Fig. 1.**
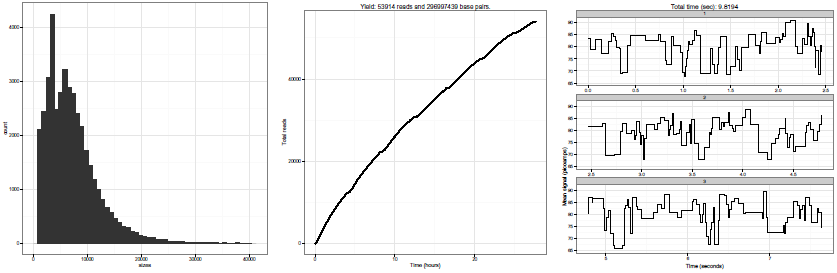
Examples of visualisations generated by `poretools` on a set of FAST5 files generated by a single MinION™ run. Panel A shows a histogram of read lengths. Panel B shows a collector’s curve of reads over time. Panel C shows a portion of an example squiggle plot of detected event transitions originating from the nanopore instrument.

As summarized in Table 1, `poretools` also provides several utilities for extracting the low level details that led to each base called sequence (see Table 1 for details). In particular, the `events` utility reports the mean current observed for each nanopore translocation event, as well as the time (in milliseconds) of each event and the k-mer that was predicted to have occupied the nanopore during the event. The `squiggle` utility permits visualization of this information (Figure 1C). The Oxford Nanopore base-calling software employs a Hidden Markov Model to predict a fragment’s sequence based upon this event data. We anticipate that the `events` utility (and others) will help new developers explore improved base-calling strategies.

**Table 1.**
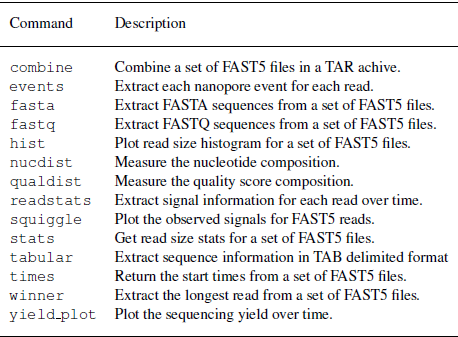
Summary of currently supported operations in poretools.

### 2.3 Python library for data analysis

The utilities provided in the `poretools` suite will inevitably prove to be insufficient for every analysis that a researcher wishes to conduct. Recognizing this, we have deveoped a Python programming interface that researchers can use to directly access the sequence data, the raw nanopore event data, and other metadata (e.g., the flowcell and and run identifiers) contained in one or more FAST5 files. In order to demonstrate the use of the Python interface, the following code reports the start time, the specific nanopore, and the based-called sequence for each FAST5 file in a given sequencing run.

~~~
**from**  poretools **import** Fast5FileSet
~~~

~~~
fast5s = Fast5FileSet (’ / path / to / fast5 / files / ‘)
**for** fast5 **in** fast5s :
  start = fast5 . get_start_time ()
  porenum = fast5 . get_channel_number ()
  fq = fast5 . get_fastq ()
  **if** fq **is not** None :
          **print** porenum , start , fq . seq , fq . qual
  fast5 . close ()
~~~

## 3 DISCUSSION

The `poretools` software helps solve pressing requirements for analysis of nanopore sequencing data. By focusing on the Python development environment and adopting expected interface conventions as popularised by other popular bioinformatics tools such as `samtools` (Li *et al*. (2010)) and `bedtools` (Quinlan *et al*. (2009)), we expect that users will be able to rapidly exploit the functionality offered by this software. We anticipate that other toolkits will become available written in other programming languages. Further efforts are required for downstream analysis for common tasks including alignment and *de novo* assembly of both event and base-called sequence data from this platform.

## ACKNOWLEDGEMENT

The authors would like to express their gratitude to ONT for granting access to the MinION™ early access programme. We thank the employees of ONT for assistance during this programme, with particular thanks to Zoe McDougall, Clive Brown, Daniel Turner, Stephanie Brooking, Roger Pettett and Stuart Reid. We would like to thank members of the MinION user community who have tested `poretools` and continue to provide feedback and bug reports.

### Funding

NJL is funded by a Medical Research Council Special Training Fellowship in Biomedical Informatics. ARQ was supported by an award from the NIH “(NGHRI; 1R01HG006693-01).

